# Combination of High Dose Rate Radiations (10X FFF/2400 MU/min/10 MV X-rays) and Paclitaxel Selectively Eliminates Melanoma Cells

**DOI:** 10.1101/2021.07.22.453100

**Authors:** Niraj Lodhi, Sreeja Sarojini, Michaela Keck, Poonam Nagpal, Yuk Ming Chiu, Zeenath Parvez, Laura Adrianzen, K. Stephen Suh

## Abstract

**Purpose:** Melanoma is one of the most aggressive cancer with 1.6% of total cancer deaths in United States. In recent years treatment options for metastatic melanoma have been improved by the FDA approval of new therapeutic agents. However, these inhibitors based therapies are non-specific and have severe toxicities including hyperkeratosis, photosensitivity, hepatitis, arthralgia and fatigue. The aim of this study is to determine the synthetic lethal effect (paclitaxel and radiations) on melanoma cells and reduce the total radiation doses by increasing the dose rates up to 2400 MU/min.

**Methods:** We previously reported a radiation treatment (10 MV x-rays, 10X-FFF, dose rate 2400MU/min, low total dose 0.5 Gy) that kills melanoma cells with 80% survival of normal HEM in vitro. In this study we extended the radiation cycle up to four and include paclitaxel treatment to study the synthetic lethal effect on melanoma and two additional normal primary cells, HDF and HEK. Cells were treated with paclitaxel prior to radiations of dose rate of 400 and 2400 MU/min with total radiation dose of only 0.5 Gy. To study induction of apoptosis and cell death, mitochondria respiration assay, DNA damage assay and colony formation assay were performed.

**Results:** Four days of consequent radiation treatment with paclitaxel significantly reduces the survival of melanoma cells by inducing of apoptosis and mitochondrial damages. After treatment, excessive DNA damage in melanoma cells leads to increase in expression of pro-apoptotic genes (Casp3) and decrease in expression of DNA repair gene (PARP1) and anti-apoptotic gene (Bcl2) to activate apoptosis pathway. Combination of paclitaxel and radiations reduces the survival of melanoma cells colonies when compared to radiation alone.

**Conclusion:** Our study indicates radiations with paclitaxel has potential synthetic lethal effect on melanoma cells and can be develop as therapy for melanoma without having toxicities or harmful effects to normal primary skin cells.

## 1. Introduction

After identification of BRAF mutation, one of the important and major drivers of melanoma, it has led to the development of targeted therapies including immunotherapy, single agent or combination chemotherapy and radiotherapy [1]. However, the ten-year survival rate for metastatic melanoma continues to be 10%, and the recent annual number of estimated newly diagnosed cases is greater than 73,000 with death numbers reaching almost 10,000 [2]. Melanoma was known to be a radio-resistant tumor initially [3] but as the radiotherapeutic measures evolved, it was proved that it is radio-sensitive and radiation therapy option is open [4]. Stereotactic body radiotherapy (SBRT) uses high dose rate of Flattening Filter Free (FFF) beams for cancer therapy because of the significantly shortened beam on time. A study of patients (n=84) with multiple lesions (lung 75, liver 10, adrenal 6, lymph nodes 5, others 4) using 6 or 10 MV FFF beams demonstrated no acute toxicity on patients with overall survival (OS) at 94%, indicating that radiotherapy can be used in melanoma treatment [5]. For unresectable malignant melanoma in the esophageal tract demonstrated that a chemo-radio combination is helpful in preventing supraclavicular metastasis with favorable palliative effects [6].

Paclitaxel (microtubules disassembly protector, blocks the progression of mitosis and triggers apoptosis) was introduced in cancer therapy decades ago and has been widely used for ovarian [7–10], breast [11, 12], Kaposi-Sarcoma (AIDS-related) [13–15] and lung carcinomas as a single agent or in combination with other drugs [16–20]. In multiple studies paclitaxel was investigated for its effectiveness to treat metastatic melanoma either as single agent or in combination with a small molecule inhibitor [21] small molecule inhibitors and carboplatin [22–24] or immunocytokine F8-IL2 [25]. For non-resection metastatic melanoma, paclitaxel is currently in clinical trial for first line therapy with other chemotherapy drugs with reporting of promising primary results [26]. Recent clinical practice uses chemo agents with radiotherapy for cancers. The paclitaxel and cisplatin combinations were used with Intensity-Modulated Radiation Therapy (IMRT) in a study in treating upper esophageal carcinoma with favorable results and no significant toxicities [27].

Here, we report the apoptotic effect on melanoma cells by combining the low dose paclitaxel with the high dose rate/low total dose (radiation) method. We treated both melanoma and normal primary skin cells Human Epidermal Melanocytes (HEM), Human Epidermal Keratinocytes (HEK) and Human Dermal Fibroblasts (HDF) for four consecutive days with a combination protocol to analyze the accumulative apoptotic effects on cancer cells in vitro. All melanoma cells were eliminated after four treatments and all normal primary skin cells were preserved and unharmed.

## 2. Materials and methods

### 2.1 Cell culture

Melanoma cell line WC00046 (V600E mutation in BRAF gene) (Figure 1A) purchased from Coriell Institute (Camden, NJ) was cultured in RPMI medium containing 10% FBS, 1% penicillin/streptomycin (Invitrogen, Grand Island, NY). HEM and culture media were purchased (ScienCell, Carlsbad, CA), and HDF and HEK were prepared as previously described [28, 29].

**Figure 1.**
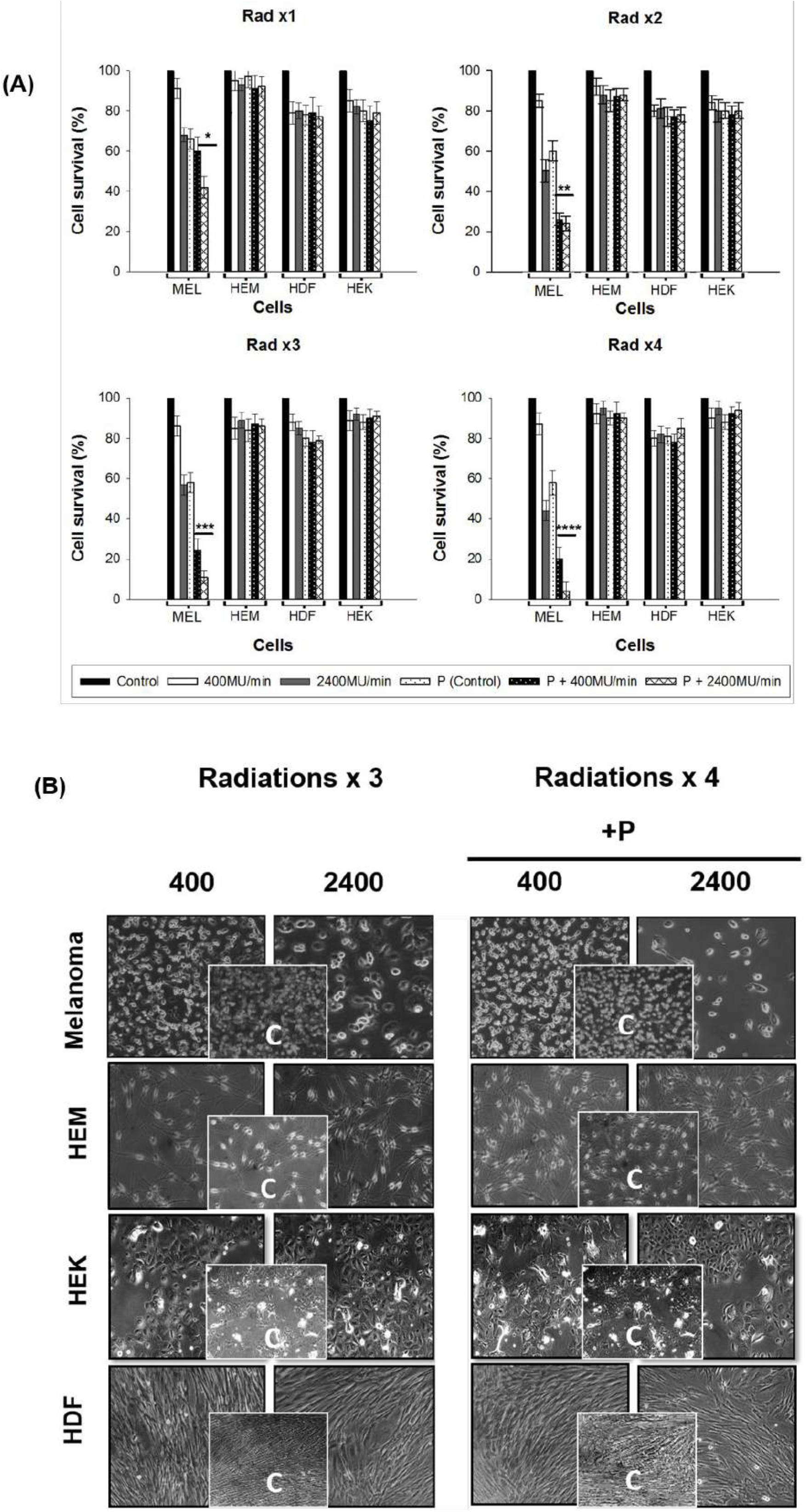
Melanoma cell line (WC00046), normal skin cells (HEM, HEK, and HDF) were irradiated (Rad x1 to x4) with and without paclitaxel. **(A).** Melanoma cell lines showed significant reduction in cell survival % after the third radiation in paclitaxel treated cells. Normal cells survival was >80% even after treating with paclitaxel. The statistical significance between the two dose rates for WC00046 with paclitaxel (*) is p<0.002, for Rad x2 is p<0.006 (**), Rad x3 is p<0.001(***), and Rad x4 p<0.002 (****). **(B). Live imaging of melanoma or normal cells irradiated cells.** Cells in T25 culture flasks using the IDEA camera, Spot 5 by phase contrast microscopy after 4 radiations (Rad x4). Control cells from the melanoma cell line, HEM, HEK and HDF are given in inset. Each cell line radiated under 400 MU and 2400MU with and without paclitaxel is visualized in this image.

### 2.2 Radiation treatments to cells

Cells were seeded (2x 105) in T-25 culture flasks (BD-Falcon), allowed to adhere overnight, and irradiated with 10 MV x-rays at dose rates of 2400 MU/min or 400MU/min by using TrueBeam (Varian Medical Systems, CA) with 10X-FFF mode. The radiation dose was administered to melanoma and normal cells as shown in Supplementary Figure 1 and Supplementary Table 1. Where a duplicate set of cells of each group was pretreated with paclitaxel (50nM in DMSO; Sigma Aldrich,A) for two hours prior to each irradiating step (Rad x1, Rad x2, Rad x3, and Rad x4). The cell culture medium was changed after two hours of each radiation. Cells were treated with paclitaxel only two hours prior to each irradiation. The titrated dose of 50nM paclitaxel was not toxic to normal skin cells for the entire duration of the experiment.

### 2.3 Colony formation assays

One day after radiation, HEK, HEF, HEM, and melanoma cells were treated with trypsin, collected, and serially diluted (1:100, 1:1000, and 1:10000) for appropriate seeding in dishes (Corning, NY) with complete media. Colonized cells (typically 21 days) were stained with hematoxylin for 30 min, fixed with 100% ethanol for 30 min, washed in water and dried overnight for counting. Radiated cells were counted with a Beckman Coulter Counter (Brea, CA).

### 2.4 RNA isolation and quantitative PCR

The TRIzol (Invitrogen, Grand Island, NY) method was used for RNA extraction, and selected genes were amplified by Q-RT-PCR with SYBR green (Qiagen, Valencia, CA) on the FAST Model 7900HT (Applied Biosystems, Carlsbad, CA). Data were analyzed using the SDS 7900HT software v2.2.2 application to determine the comparative threshold cycle (Ct) method (2-ΔΔCt) for calculating fold changes and standard deviations [30]. Primer sequences are given in Supplementary Table 2.

### 2.5 Cell proliferation assay

The MTS assay (CellTiter 96^®^ AQueous One Solution, Promega, Madison,WI) was used to assess cell proliferation of radiated cells by using a Microplate reader (BioTek Synergy HT) as described by the manufacturer and based on previous publication [28].

### 2.6 Mitochondria respiration assay

Mitotracker Red CMXRos (Invitrogen-M7512) was used to stain cells at a final concentration of 200nM for analyzing mitochondria activity before and after radiation. After a 15-minute incubation at 37°C and 5% CO2, cells were washed with PBS and fixed with 2% PFA (paraformaldehyde) for 30 minutes at room temperature in the dark. Fixed cells were washed with PBS, mounted using DAPI, and fluorescent images were taken immediately with Zeiss Fluorescent microscopy (Axiovision). Cell fluorescence was quantified by using ImageJ.

### 2.7 DNA damage assay

The EpiQuik In Situ Kit (Cat # P-6001-096, Epigentek, Farmingdale NY) detected the phosphorylation of H2AX at Serine 139 by a colorimetric method measured at 450nm in a 96-well plate where the cells are grown after radiation, fixed, and permeabilized according to the manufacturer’s protocol. The DNA damage assay was carried out one hour after paclitaxel and the radiation treatment.

### 2.8 Western blotting

Western blotting experiments were carried out by using cell lysates (Pierce IP lysis buffer cat# 87788; Invitrogen) prepared from cell lines treated with or without Paclitaxel or radiation x4. SDS-PAGE gels (12%, Criterion TGX Precast Gels) were used to fractionate cell lysates and transferred onto a polyvinylidene difluoride (PVDF, Bio Rad) membrane using a Bio-Rad transfer unit (Hercules, CA). The membranes were blocked by using 5% nonfat dry milk in TBST (10 mmol/l Tris, pH 8.0, 150 mmol/l NaCl, 0.5% Tween 20) for 60 min; they were then washed and incubated with primary antibodies against Bcl-2 (1: 1000, #7973; Abcam, Cambridge, MA), caspase-3 (1:2000, H-277), PARP 1 (1:2000, #SC-7148, SC-8007; Santa Cruz Biotechnology Inc., Santa Cruz, CA) and Actin (1:10 000, AB-6276) at 4°C overnight. Actin was used as the loading control. The membranes were then washed and incubated with a 1:5000 dilution of horseradish peroxidase-conjugated donkey anti-mouse IgGs secondary antibody. The blots were washed and developed using the ECL system (Pierce). Doxorubicin (Dx, topoisomerase II inhibitor) treated cell lysates were used as a positive control (+C) for Caspase3 and negative control for PARP1 and Bcl-2. Samples were used from a Rad x4 experimental setting with (+Pac) or without (-Pac) Paclitaxel and with (T) or without (U) radiation treatment.

### 2.9 Statistical analysis

All experiments were performed a minimum of three times, and data represent the results for assays performed in triplicate or quadruplicate. Error bars represent 95% confidence intervals (CIs). All statistics were based on continuous variables by using software STAT View, and P values <0.05 were considered statistically significant. For comparisons between two groups, the Student’s t test was applied.

## 3 Results

### 3.1 Paclitaxel in combination with a high dose rate of 2400MU/min (total dose 0.5 Gy) significantly increases apoptosis in melanoma cells

Viable cells were counted four days after four consecutive days of the treatment (Figure 1A; Rad x1, x2, x3, and x4). At the high dose rate (2400MU/min total dose 2Gy Rad x4) in the presence of paclitaxel the normal skin cells tolerated the total dose of 2Gy (0.5Gy x4=2Gy) for both 400MU/min and 2400MU/min at the high dose rate, but the melanoma cells were selectively killed: 58% on day 1 (Rad x1), 76% on day 2 (Rad x2), 96% on day 3 (Rad x3), and 98% on day 4 (Rad x4) (Figure 1A, P + 2400MU/min, p<0.005 for all 4 treatments). The cell counts after the four consecutive treatments showed that melanoma cell survival percentages decreased 5-fold compared to the control (non-radiated, no paclitaxel) for P+400MU/min and 50-fold for P+2400MU/min, suggesting that 2400MU/min in combination with paclitaxel is highly effective for killing melanoma cells while largely preserving normal skin cells (Figure 1A).

Based on bright field microscopy data, all normal skin cells HEM, HDF and HEK maintained above 85% cell survival after four consecutive daily radiations with or without paclitaxel and with 400MU/min or 2400MU/min dose rate (Figure 1B), suggesting that the titrated dose of paclitaxel and 0.5Gy of total dose delivery leads to minimal harm to normal skin cells. All skin cells retained proliferative potential and recovered fully three days after the fourth treatment.

### 3.2 Paclitaxel in combination with 2400MU/min (total dose 0.5Gy) induces significantly higher DNA damage to melanoma cells

The treated cells were harvested two days after four consecutive treatments (Supplementary Table 1) and allowed to recover for at least 24 hours after each treatment prior to the subsequent radiation. Irradiating melanoma cells with 400MU/min did not increase DNA damage, but the 2400MU/min dose rate increased to three-fold more DNA damage than the control and 400MU/min (Figure 2A). Normal skin cells showed moderate DNA damage in cells with radiation alone, but less than paclitaxel treated cases (Figure 2A), Despite DNA damage, cell proliferation markers CCND1 and CCND2 were moderately upregulated in treated normal skin cells, but downregulated in treated melanoma cells (Figure 2B). The level of radioprotection gene SOD2 was downregulated in melanoma cells while HEM upregulated the expression levels, suggesting that the DNA damage repair and radioprotection of normal skin cells are actively enhancing the survival of normal cells.

**Figure 2.**
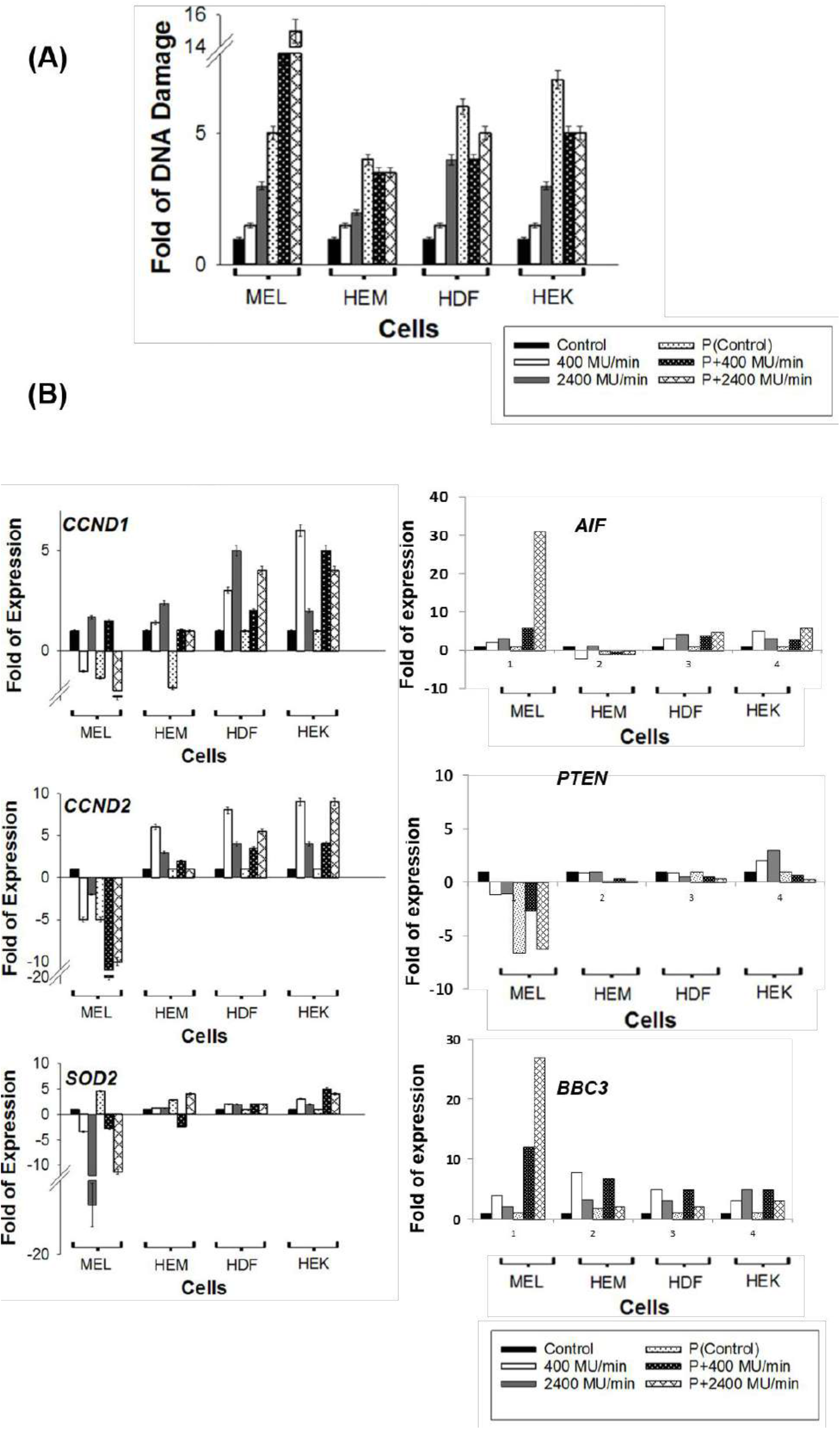
**(A).** DNA damage (fold) of irradiated cells with and without co-administration of paclitaxel for both dose rates 400MU/min and 2400MU/min are shown after the fourth radiation (Rad x4). Data represent the average of four independent experiments with error bars. The statistical difference between 400MU/min and 2400MU/min with paclitaxel is shown (**) p<0.004. **(B).** Mutational status of cell cycle genes CCND1, CCND2, Radioprotection gene SOD2, AIF, PTEN, and BBC3 were quantified via qRT-PCR analysis.

### 3.3 Proliferation of melanoma cells reduce and mitochondrial respiration increase in paclitaxel and 2400MU/min (total dose 0.5Gy) radiations treatment

MTT assay was used to investigate cell proliferation and mitochondrial respiration of normal skin cells and melanoma cells. The cell proliferation assay using MTS reagent suggested all normal skin cells HEM, HDF and HEK showed proliferation level close to or above the untreated control after four radiations, while proliferation was significantly reduced in melanoma cells after four radiations. Combination treatment is effective to reduce viability of melanoma cells – 44% (Rad x1), 31% (Rad x2), 4% (Rad x3), and 0.5% (Rad x4) versus viability of >95% after four radiations with or without paclitaxel at dose rate of 2400MU/min for normal skin cells (Figure 3A).

**Figure 3.**
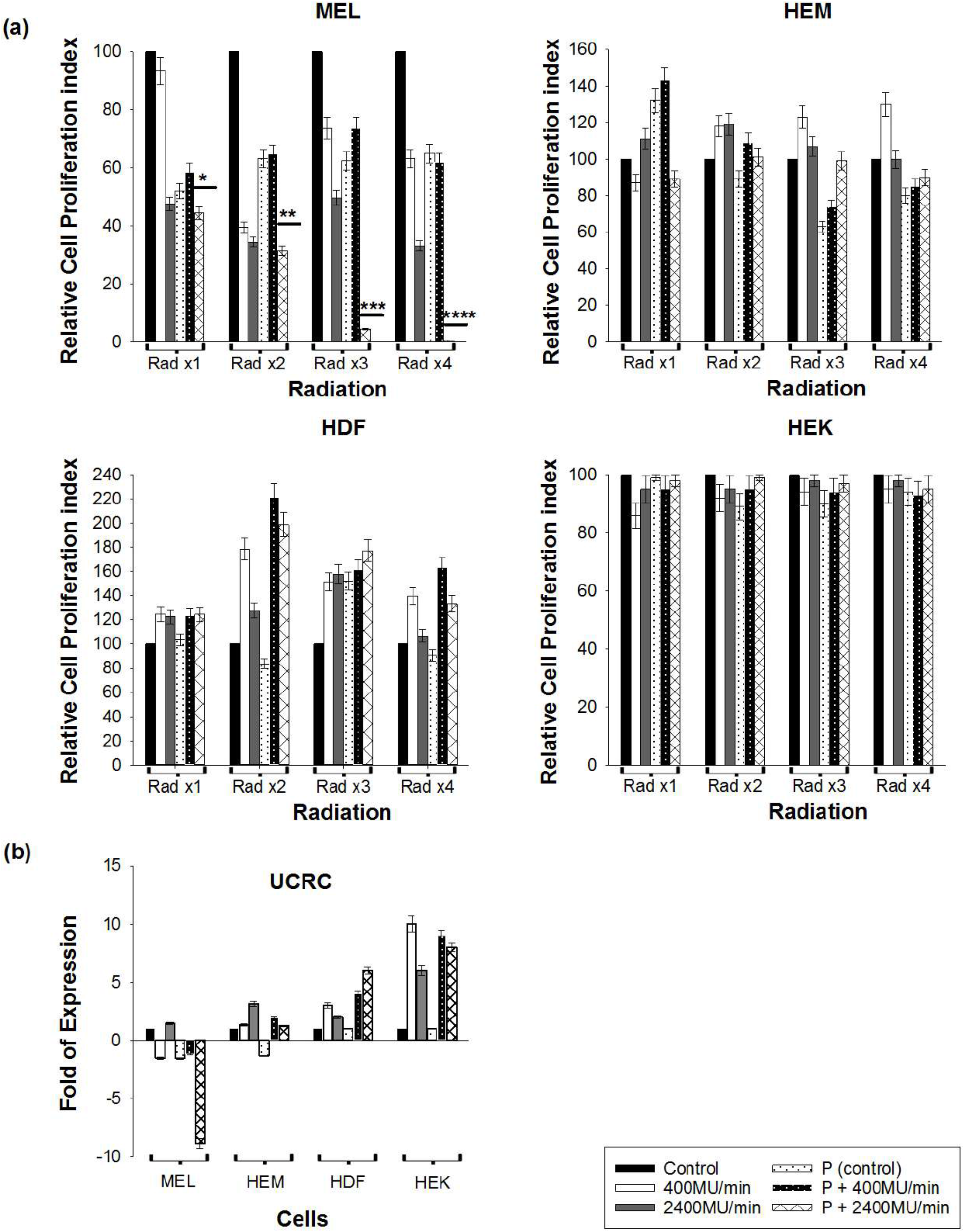
Cell proliferation was quantified by MTT assay. **(A).** Cell proliferation was quantified by MTT assay for irradiated cells from (A) and standard error bars and statistical p values are shown as (*). For the melanoma cell line, the statistical significance between 400MU/min and 2400MU/min with P (*) is p<0.005, for Rad x2 is p<0.006 (**), Rad x3 is p<0.002(***), and Rad x4 p<0.005 (****). (B) The mutational status of the mitochondrial respiration gene UCRC was analyzed using qRT-PCR, and the average fold changes against non-radiated control cells are shown.

Dose rate 2400 MU/min caused higher mitochondrial respiration than 400 MU/min which was directly related to total dose of radiation delivered to cells. However, the respiratory chain gene (UCRC) was down regulated in melanoma cells, suggesting that the increased respiration activity after radiation was related to post-translational activation (Figure 3B).

### 3.4 Expression of pro-apoptotic gene is elevated and DNA repair PARP1 gene and anti-apoptotic gene are down regulated in melanoma cells caused enhanced apoptosis

Anti-apoptotic Bcl-2 was downregulated in irradiated and paclitaxel-treated melanoma cells on the RNA and protein level (Figure 4A). Pro-apoptotic Casp3 is upregulated in irradiated and paclitaxel-treated melanoma cells on the RNA level, but showed downregulation on the protein level (Figure 4B). However, normal skin cells retain expression of Casp3 and Bcl-2 in treated and untreated cells (Figure 4A, B). DNA damage repair genes PARP1 (Figure 4C) was downregulated substantially in paclitaxel treated irradiated melanoma cells, while the expression levels were only slightly changed in normal skin cells. Protein analysis by anti-PARP1 showed the downregulation of levels in irradiated and paclitaxel-treated melanoma cells (Figure 4C), whereas untreated cells maintained base level expression.

**Figure 4.**
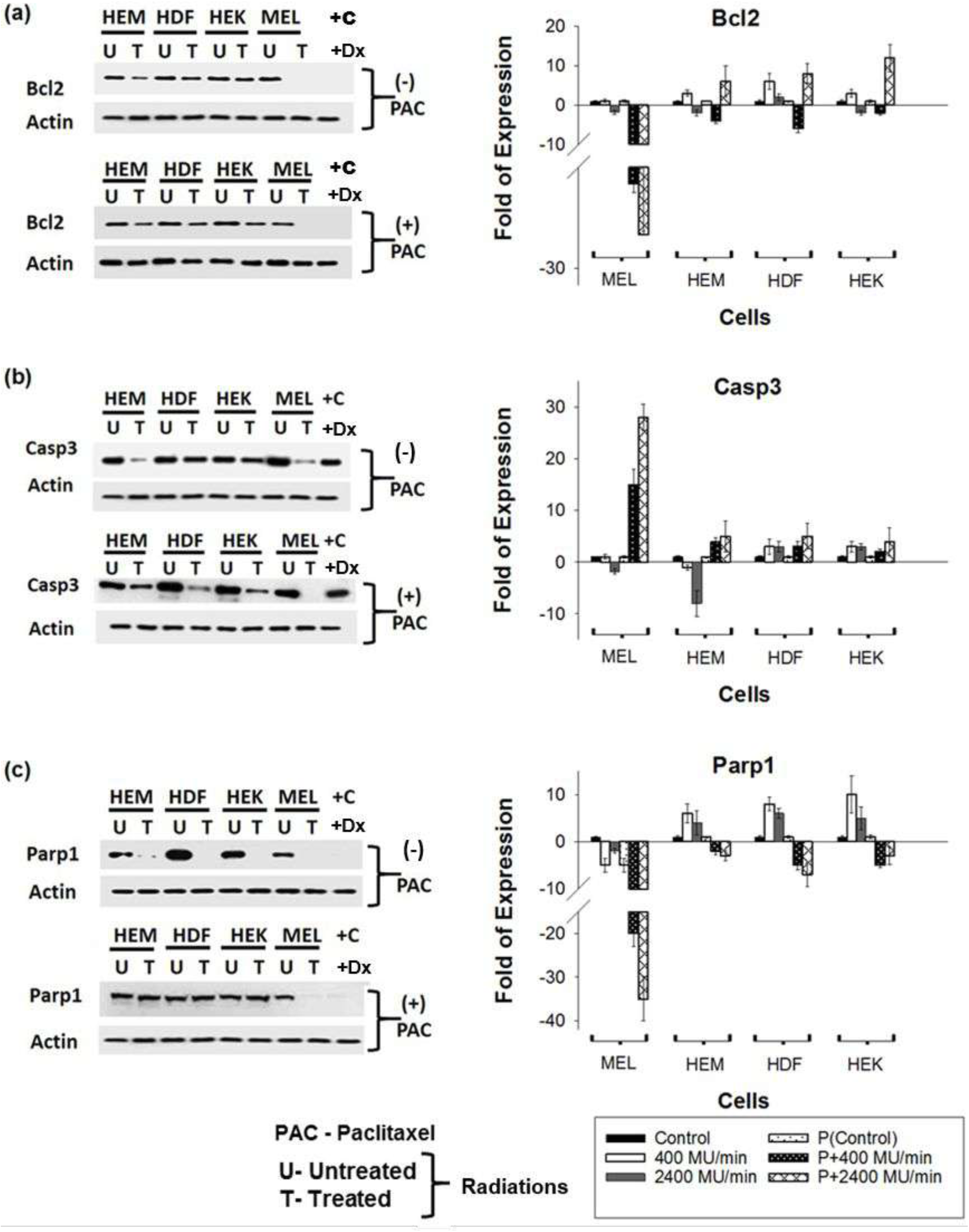
Effect of radiations and paclitaxel on the expression of DNA repair and ant-apoptosis genes. Western Blot analyses of Bcl-2 (A), Caspase3 (B), PARP1 (C), and Doxorubicin (Dx) treated cell lysates were used as positive controls (+C). The samples used were from a Rad x4 experimental setting with (+PAC) or without (-PAC) paclitaxel and with (T) or without (U) radiation treatment. Subsequently, q RT-PCR analysis of the same Bcl-2, PARP1, and Caspase 3 were also carried out, and fold expressions were shown in graphs.

A Mitotracker assay of melanoma cells treated with paclitaxel and irradiated at 2400MU/min (Rad x4) showed no fluorescence, indicating cells undergoing apoptosis (Figure 5A and B). Minimal fluorescence was noticed in cells irradiated at 400MU/min with paclitaxel even after four radiations, suggesting that paclitaxel induced more mitochondrial damage when co-administered with dose rate 2400MU/min in cancer cells HEM, HDF and HEK showed no significant difference in fluorescence between 400MU/min and 2400MU/min.

**Figure 5.**
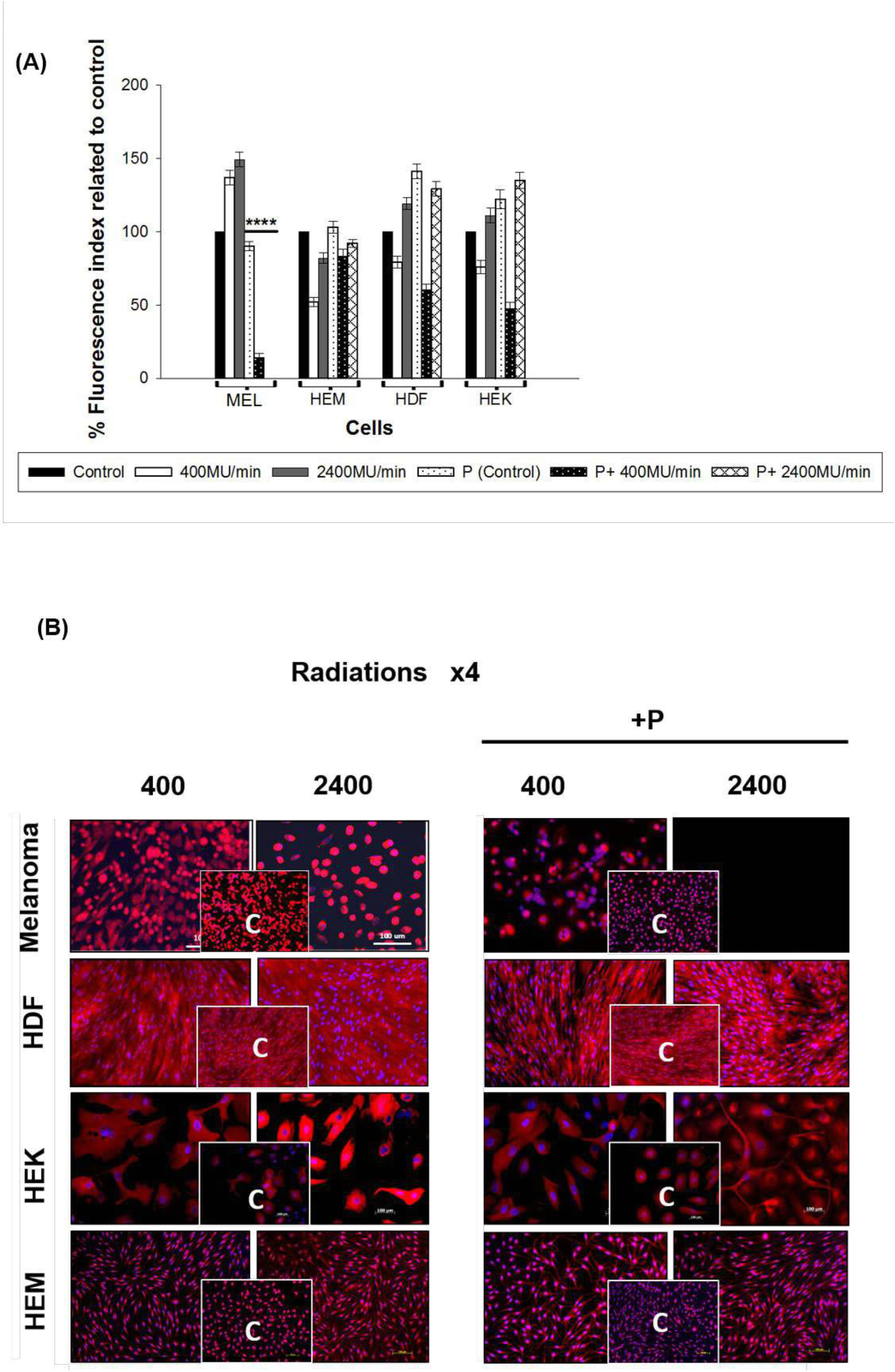
Effect of radiations and paclitaxel on mitochondrial respiration using Mitotracker Red-fluorescent dye. **(A).** The average fluorescent intensity from five random fields for each experimental setting was used to calculate the relative fluorescent intensity using Image J, and was normalized against the average intensity of the non-radiated control (solid black) for each cell type. The fold changes for dose rates 400MU/min (solid white) and 2400MU/min (solid gray) are shown with standard error bars. Paclitaxel control (P) data are represented with black dots over white; 400MU/min + P are shown in white over black, and 2400MU/min is in horizontal diamond bars. The statistical significance between the two dose rates for WC00046 with paclitaxel (*) is p<0.005). **(B).** The melanoma cell line, HEM, HDF, and HEK cells were radiated with and without paclitaxel four times (Rad x1, Rad x2, Rad x3, and Rad x4). The image represents Rad x4 data. Cells were collected and seeded and stained the following day using Mitotracker Red-fluorescent dye to detect mitochondrial respiration, and florescence microscopy was used for imaging at 20X magnification with scale bars. Non-radiated control cells (inset) are shown for individual radiation settings for each cell type.

### 3.5 Colony formation of melanoma cells diminished after four cycles of radiations at dose rate 2400MU/min (total dose 2 Gy) and paclitaxel treatment

Colony formation assay has been used as the classic protocol in radiation research for determining the ability of a single cell to multiply into a colony [32]. Data showed that 2400MU has higher apoptotic potential on melanoma cells than 400MU/min (Figure 6A). Co-administration of paclitaxel reduced the survival of colonies to 30% (Rad x1), 18% (Rad x2), 2% (Rad x3), and 0% (Rad x4) when compared to 2400MU/min alone (Figure 6A and B). For normal skin cells, there was no significant difference between survival of colonies of irradiated and paclitaxel-treated cells when compared to radiation alone.

**Figure 6.**
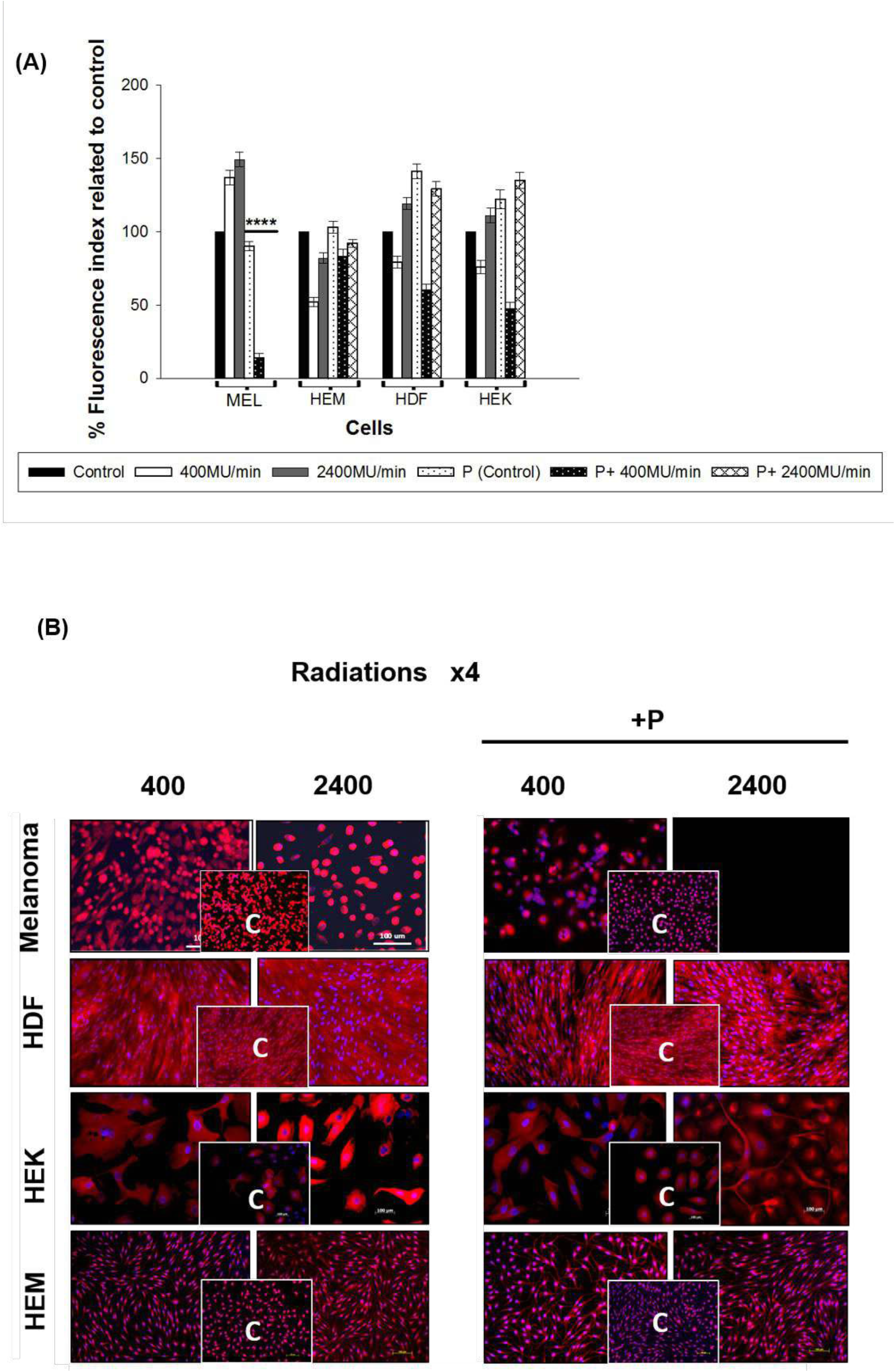
Colony formation assays after each radiation and paclitaxel experiment. **(A).** Cells were counted, serially diluted, and plated on culture grade petri dishes. After 2-3 weeks colonies were stained and counted, and the survival percent (%) was calculated. The statistical significance between the two dose rates for WC00046 with paclitaxel (*) is p<0.001, for Rad x2 is p<0.004(**), Rad x3 is p<0.005(***), and Rad x4 p<0.002 (****). **(B).** Images of colonies from the colony assay plates after staining with hemotoxylin. Melanoma cell lines were radiated with and without paclitaxel. The image represents Rad x4 data.

## 4. Discussion

Melanoma is known to be radio-resistant, and radiotherapy is currently used for palliative, but not curative care. In recent years, many scientific efforts have been made to compromise DNA repair efficiency of melanoma cells, including radiation dose. The current study demonstrates the effectiveness of combining dose rate 2400MU/min (at a low total dose of 0.5Gy) with paclitaxel, a cost-effective chemotherapeutic drug. Food and Drug Administration (FDA) approved paclitaxel induces DNA damage and mitochondrial disruptions along with microtubule inhibition [33]. We show that melanoma cell’s survival is significantly reduced after four cycles of paclitaxel and radiation treatment at dose rate of 2400MU/min. while minimal affect in normal skin cells HEM, HDF and HEK (Figure 1). We measured DNA damage caused by radiations by phosphorylation of H2AXSer139 as was used by other research groups [34–41] in primary or melanoma cells. This assay shows that radiations at 2400MU/min with paclitaxel induce prolonged DNA damage in melanoma cells and cause six-fold more DNA damage than 2400MU/min alone (Figure 2A).

Excitation of mitochondria after exposure to ionizing radiation is related to upregulation of mitochondrial electron transport chain function which helps the cancer cells to become resistant to radiation [42]. Paclitaxel has the ability to damage the mitochondria by altering mitochondrial respiration and thus induce mitochondrial apoptosis [43]. We used MitoTracker probes, which passively diffuse across the plasma membrane and accumulate in active mitochondria in viable cells only [44–46]. Further, to confirm induction of DNA Damage Response (DDR) and pro or anti-apoptotic proteins to radiation exposure we performed western as in Figure 4, this combination upregulates AIF, BBC3, PARP1, and Casp3 genes in cancer cells to >25 fold. Cell cycle genes (CCND1/CCND2, Figure 2), DNA damage repair genes (PARP1, Figure 4), radioprotection genes (SOD2, Figure 2), and anti-apoptotic genes (Bcl2, Figure 4) were upregulated in normal skin cells while mitochondrial and extrinsic apoptotic pathway genes (AIF and BBC3) were upregulated in paclitaxel-treated, irradiated cells.

The colony formation assay assesses the cell reproductive death after radiation that is determined by the proportion of surviving colonies for each dose administered [47]. Our data show the decrease in cell survival for melanoma cells after paclitaxel treatment. Results were confirmed with the MTS assay, where a decrease in viability of melanoma cells was recorded when combined with paclitaxel, while normal cells retained their viability >95% (Figure 3A).

When melanoma cells are concurrently treated with paclitaxel, cell number and mitochondrial efficiency decrease and cells show low fluorescence for 400MU/min and no fluorescence for 2400MU/min after four radiations. This suggests that paclitaxel indues greater mitochondrial damage when co-administered with a dose rate of 2400MU/min in melanoma cells. As shown in Figure 5 A and B viability of paclitaxel and radiation exposure cells is very low or none.

Co-administration of paclitaxel completely inhibited the survival of colonies in melanoma cells after four radiations at dose rate of 2400MU/min (Figure 6). Similar to the standard radiotherapy (400MU/min), melanoma cells exhibit resistance to 400MU/Min in Rad x1 to Rad x4 (total dose 2Gy), even in the presence of paclitaxel; but our studies suggest that high dose rate radiation (2400MU/min of total dose 2Gy) completely abolishes the melanoma cells in combination with paclitaxel. In other studies paclitaxel was used with chemo and radiotherapy [31, 48]. These studies are in clinical trials phase and their primary outcome measures are to improve the Overall Response Rate (ORR), Proliferation Free State (PFS) or Overall Survival (OS). Paclitaxel-based chemo-radiotherapy for non-small cell lung cancer (NSCLC) patients demonstrated that minimum toxicity and an optimal radio-chemo combination regimen are yet to be established with further studies [31]. Paclitaxel was also used in melanoma frontline therapy as single agent or in combination with carboplatin (DNA damaging agent) immunotherapeutic drugs (Ipilimumab) in a phase II clinical trial to find a new approach to increase survival rates of patients [48].

Our study is new approach for radiotherapy by increasing the dose rate and reduce the total dose (0.5 Gy) to lower toxic effects of radiations in combination of paclitaxel for treatment of BRAF mutant patients. In general patients get the 40-60 Gy of radiation dose in 4-5 cycles and that cause the severe side effects of other skin problems, fatigue and radiation pneumonitis. Although this is in vitro study but death of melanoma cells is significant and no toxic effects on primary cells at total dose of 2 Gy in four cycles.

In conclusion, our study is a significant finding that supports the use of radiotherapy with chemo for melanoma when standard radiotherapy fails to kill melanoma cells due to radio-resistance. Although future studies using in vivo models will be required to prove this concept for clinical settings, this study indicates the effectiveness of paclitaxel when combined with a high dose rate of radiation (2400MU/min) and a low total dose (0.5Gy) in melanoma therapy.

## Supporting information

Radiation dose administration to melanoma or normal cells.

Cell groups for Paclitaxel or Paclitaxel with irradiation treatment.

Primer sequences used in this study.

## Acknowledgements

We thank The John Theurer Cancer Center and Radiology and Oncology Department of the Hackensack University Medical Center. We are thankful to the Lisa B. Fishman Foundation and the John Theurer Cancer Center of the Hackensack Meridian Health network for continuous support during preparation of this manuscript. We thank Esra Uckun Kiran and Shermineh Bradford for participating in Radiobiology experiments.

## Conflict of Interest Statement

No potential conflicts of interest were disclosed by the authors.

## SUPPLEMENTAL FIGURE AND TABLES

**Supplementary Figure 1.**

**Radiation dose administration to melanoma or normal cells.**

Cell groups were divided into one radiation (Rad x1, on day 1), second radiation (Rad x2; day 2), third radiation (Rad x3; day 3), and fourth radiation (Rad x4; day 4).

**Supplementary Table 1.**

Cell groups for Paclitaxel or Paclitaxel with irradiation treatment.

**Supplementary Table 2.**

Primer sequences used in this study.

## Notes

### Competing Interest Statement

The authors have declared no competing interest.

