## Supplementary figures and images for "Combination of High Dose Rate Radiations (10X FFF/2400 MU/min/10 MV X-rays) and Paclitaxel Selectively Eliminates Melanoma Cells"

### Radiation dose administration to melanoma or normal cells.

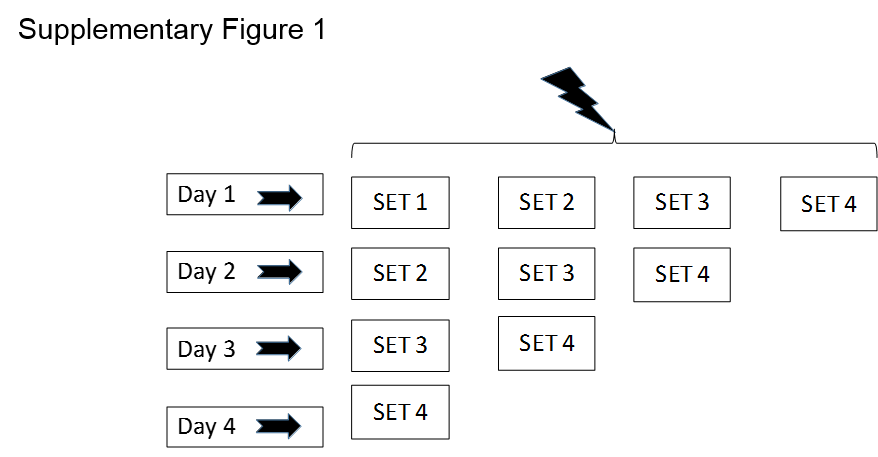
