## Supplementary material for "Combination of High Dose Rate Radiations (10X FFF/2400 MU/min/10 MV X-rays) and Paclitaxel Selectively Eliminates Melanoma Cells": Cell groups for Paclitaxel or Paclitaxel with irradiation treatment.

**Table 1. Cell groups for Paclitaxel or Paclitaxel with irradiation treatment**

|  | Control | Paclitaxel (50nM) | 400MU/0.5Gy | 400MU/0.5Gy  + Paclitaxel (50nM) | 2400MU/0.5Gy | 2400MU/0.5Gy  + Paclitaxel (50nM) |
| --- | --- | --- | --- | --- | --- | --- |
| Melanoma cells | 1*4 | 1*4 | 1*4 | 1*4 | 1*4 | 1*4 |
| HEM | 1*4 | 1*4 | 1*4 | 1*4 | 1*4 | 1*4 |
| HDF | 1*4 | 1*4 | 1*4 | 1*4 | 1*4 | 1*4 |
| HEK | 1*4 | 1*4 | 1*4 | 1*4 | 1*4 | 1*4 |
