## Supplementary material for "Combination of High Dose Rate Radiations (10X FFF/2400 MU/min/10 MV X-rays) and Paclitaxel Selectively Eliminates Melanoma Cells": Primer sequences used in this study.

| **Primer sequences used in this study** | |
| --- | --- |
| Casp3 f | gaactggactgtggcattga |
| Casp3 r | tcaagcttgtcggcatactg |
| Bcl-2 f | ttccagagacatcagcatgg |
| Bcl-2 r | tgtccctaccaaccagaagg |
| PARP 1 f | gctcctgaacaatgcagaca |
| PARP 1 r | tcctgatgatctcggcttct |
| SOD2 f | gggagatgttacagcccagata |
| SOD2 r | agtcacgtttgatggcttcc |
| UCRC f | attcgctgttggcaagaaac |
| UCRC r | tttgcagagggctttgaagt |
| PTEN f | gaatggagggaatgctcaga |
| PTEN r | cgcaaacaacaagcagtgac |
| CCND1 f | ctctcattcgggatgattgg |
| CCND1 r | gtgagctggcttcattgaga |
| GAPDH f | tcaccagggctgcttttaac |
| GAPDH r | atgacaagcttcccgttctc |
| CCND2 f | tgcagaaggacatccaaccc |
| CCND2 r | gccaagaaacggtccaggta |

**Supplementary Table 2**
